# Neurogenin regulates effectors of migratory neuron cell behaviors in *Ciona*

**DOI:** 10.1101/654798

**Authors:** Susanne Gibboney, Kwantae Kim, Florian Razy-Krajka, Wei Wang, Alberto Stolfi

## Abstract

The bipolar tail neurons (BTNs) of *Ciona* develop according to a highly dynamic, yet highly stereotyped developmental program and thus could serve as an accessible model system for neuronal delamination, migration, and polarized axon outgrowth. Here we used FACS/RNAseq to profile the transcriptional output of Neurogenin in the BTNs, searching for candidate effectors of BTN cell behaviors. We identified several candidate genes that might play conserved roles in similar cell behaviors in other animals, including mammals. Among the more interesting candidates were several microtubule-binding proteins and TGFβ pathway antagonists. A small Gαi subunit was also found to be upregulated in migrating BTNs, and interfering with its function through expression of a dominant negative inhibited delamination and a complete epithelial-to-mesenchymal transition. We propose models for the regulation of BTN behaviors by the identified candidate effectors, establishing a foundation for testing effector gene functions that might be conserved in chordate neurodevelopment.

## Introduction

In spite of an emerging picture of the molecular mechanisms of cellular morphogenesis in neurodevelopment, it is not well understood how these pathways are regulated in different developmental contexts The simple embryos of the invertebrate chordate *Ciona* are tractable for high-resolution functional genomics (Horie et al., 2018; Racioppi et al., 2019; Reeves et al., 2017; Wang et al., 2019) and *in vivo* imaging (Bernadskaya et al., 2019; Cota and Davidson, 2015; Hashimoto et al., 2015; Mizotani et al., 2018; Veeman and Reeves, 2015), and have been increasingly used to investigate the regulation of cell behaviors in development (Bernadskaya and Christiaen, 2016). Furthermore, their classification in the tunicates, the sister group to the vertebrates (Delsuc et al., 2006), means they share with vertebrates many chordate-specific gene families, cell types, organs, and developmental processes (Abitua et al., 2015; Christiaen et al., 2002; Dufour et al., 2006; Ermak, 1977; Hervé et al., 2005; Kugler et al., 2008; Ogasawara and Satoh, 1998; Razy-Krajka et al., 2012; Stolfi et al., 2010; Stolfi et al., 2015; Stolfi et al., 2011; Tolkin and Christiaen, 2012), particularly their larval central nervous system (CNS), a miniaturized but typically chordate CNS containing only 177 neurons (**Fig 1a**)(Ryan et al., 2016). *Ciona* are thus model organisms well-suited to the study of potentially conserved, chordate-specific gene regulatory networks controlling cell behaviors during the neurodevelopment.

**Figure 1.**
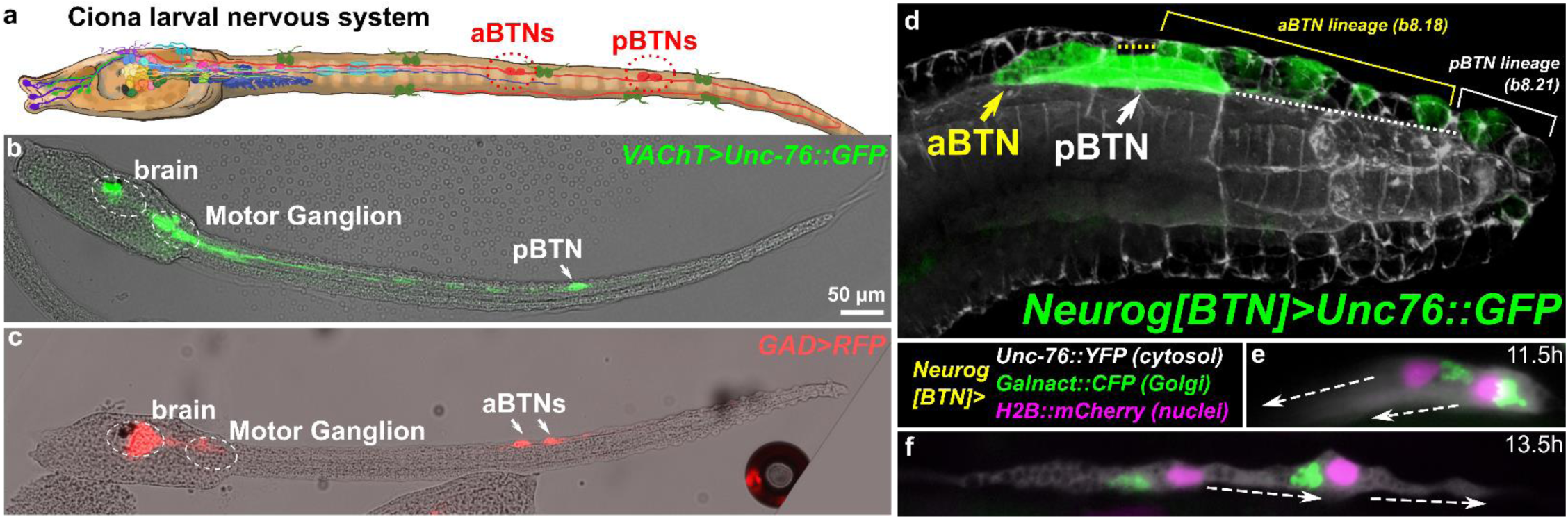
Ciona Bipolar Tail Neurons and the larval nervous system. a) Cartoon diagram of Ciona larval nervous system based on (Ryan et al., 2016), showing approximate positions of posterior BTNs (pBTN) and anterior BTNs (aBTNs). b) Cholinergic (Kratsios et al., 2012) and c) GABAergic (Zega et al., 2008) reporters label pBTNs and aBTNs respectively. Larva in (b) appears to be a left/right mosaic, which is why only 1 pBTN is visible. d) Confocal image of migrating BTNs in tail tip of a tailbud embryo electroporated with *Neurog[BTN]>Unc-76::GFP* (green), e) Relative position of Golgi apparatus is posterior to the nucleus in the BTNs during their migration forward, then f) becomes anterior to each nucleus during distal process extension. Larva diagram illustration by Lindsey Leigh.

Two of the earliest cell behaviors that are crucial for neuronal function and final connectivity are polarization and migration. Neurons develop highly polarized morphologies that are key to their function and connectivity. Various mechanisms contributing to neuronal polarization have been elucidated in different systems (Polleux and Snider, 2010; Yogev and Shen, 2017), but there is ongoing debate on the relative contributions of *intrinsic* and *extrinsic* signals in neuronal polarization *in vivo*. In the descending decussating neurons of the *Ciona* Motor Ganglion (MG), we identified the centrosomal microtubule-bundling/stabilizing protein Nckap5 as being potentially being required for proper axon outgrowth (Gibboney et al., 2019). Another centrosome-enriched protein of unknown function, Efcab6-related, and the extracellular axon guidance cue Netrin1 are also hypothesized to be required for ddN axon outgrowth (Gibboney et al., 2019), though the expression of Netrin1 by the ddNs themselves blurs the distinction between *intrinsic/extrinsic* signals, suggesting that autocrine secretion of extracellular signals by the neuron could be an essential component of an intrinsic mechanism for its polarization.

To further study the regulation of these cell behaviors in *Ciona* neurons, we focused on the Bipolar Tail Neurons (BTNs, **Fig 1b,c**). The BTNs are two bilateral pairs of neurons located along the tail nerve cord and derive their name from the two long processes they extend in opposite directions along the anterior-posterior axis. Each left/right pair is comprised of an anterior BTN (aBTN) and a posterior BTN (pBTN) that arise from separate but adjacent lineages (**Fig 1d**). The BTNs are proposed homologs of vertebrate dorsal root ganglia (DRG) neurons, based on their developmental origin from the neural crest-like cells, their early expression of Neurogenin (Neurog) family of proneural transcription factors, their morphogenesis, and their role in relaying peripheral sensory information to the CNS (Stolfi et al., 2015). Like neural crest-derived DRG neurons in vertebrates, BTNs delaminate from the dorsal midline ectoderm and migrate along somitic mesoderm as a simple chain comprised of the aBTN followed by the pBTN (**Fig 1d**), achieving their unique morphology by first extending a neurite anteriorly (**Fig 1e**), then reversing polarity and extending a neurite posteriorly (**Fig 1f**). Sustained expression of Neurog is necessary and sufficient for BTN specification, as supernumerary BTNs generated by ectopic Neurog overexpression engage in these same stereotyped behaviors (Stolfi et al., 2015). In vertebrates, Neurog1/Neurog2 are also expressed during the specification of DRG neurons as they migrate through somatic mesoderm and begin to differentiate into their bipolar (more accurately pseudounipolar) shape to transmit sensory information from peripheral tissues to the CNS (Ma et al., 1999). Therefore, Neurog factors might be activating conserved regulatory “programs” for neuronal migration, polarization, and axon outgrowth that are shared between tunicates and vertebrates.

Here we report the use of RNAseq on migrating BTNs under Neurog gain- or loss-of-function conditions, dissociated and isolated from synchronized embryos using fluorescence-activated cell sorting (FACS). By analyzing BTN transcriptome profiles under these conditions, we identified, and validated by *in situ* hybridization, a core set of candidate “effector” genes downstream of Neurog. These genes encode a diverse set of intracellular and extracellular proteins that have begun to paint a picture of the molecular pathways that might be important for BTN delamination, migration, and morphogenesis. By testing one such candidate, a small Gα_i_–like protein subunit that is specifically upregulated in the BTNs and notochord, we have identified a potential developmental switch to induce cell motility. This study sets a foundation for the dissection of a potentially conserved, chordate-specific transcriptional network for developing sensory neurons, and for further understanding the developmental regulation of morphogenetic cell behaviors.

## Results and discussion

### RNAseq profiling of potential Neurog targets in isolated BTN progenitors

Using FACS-RNAseq (**Fig 2a**) we profiled cells labeled with a *Neurog[BTN]* fluorescent reporter under different experimental conditions, isolated from synchronized embryos at 9.5 hours post-fertilization (hpf) at 20°C. In the “control” condition (*Neurog>lacZ*) only 4 cells per embryo become BTNs, while the rest of the BTN lineage is initially specified as broadly epidermis (~15-16 cells at mid-tailbud), with various epidermal sensory neurons specified later (**Fig 2b**)(Stolfi et al., 2015). In parallel, we sorted cells from embryos in which wild-type Neurog was overexpressed (*Neurog>Neurog*), or a dominant-repressor form of Neurog (*Neurog>Neurog::WRPW*). *Neurog>Neurog* specifies all cells as supernumerary BTNs, while *Neurog>Neurog::WRPW* abolishes BTN fate (**Fig 2b**). cDNA libraries were prepared from isolated cells, with each condition represented by two biologically independent replicates.

**Figure 2.**
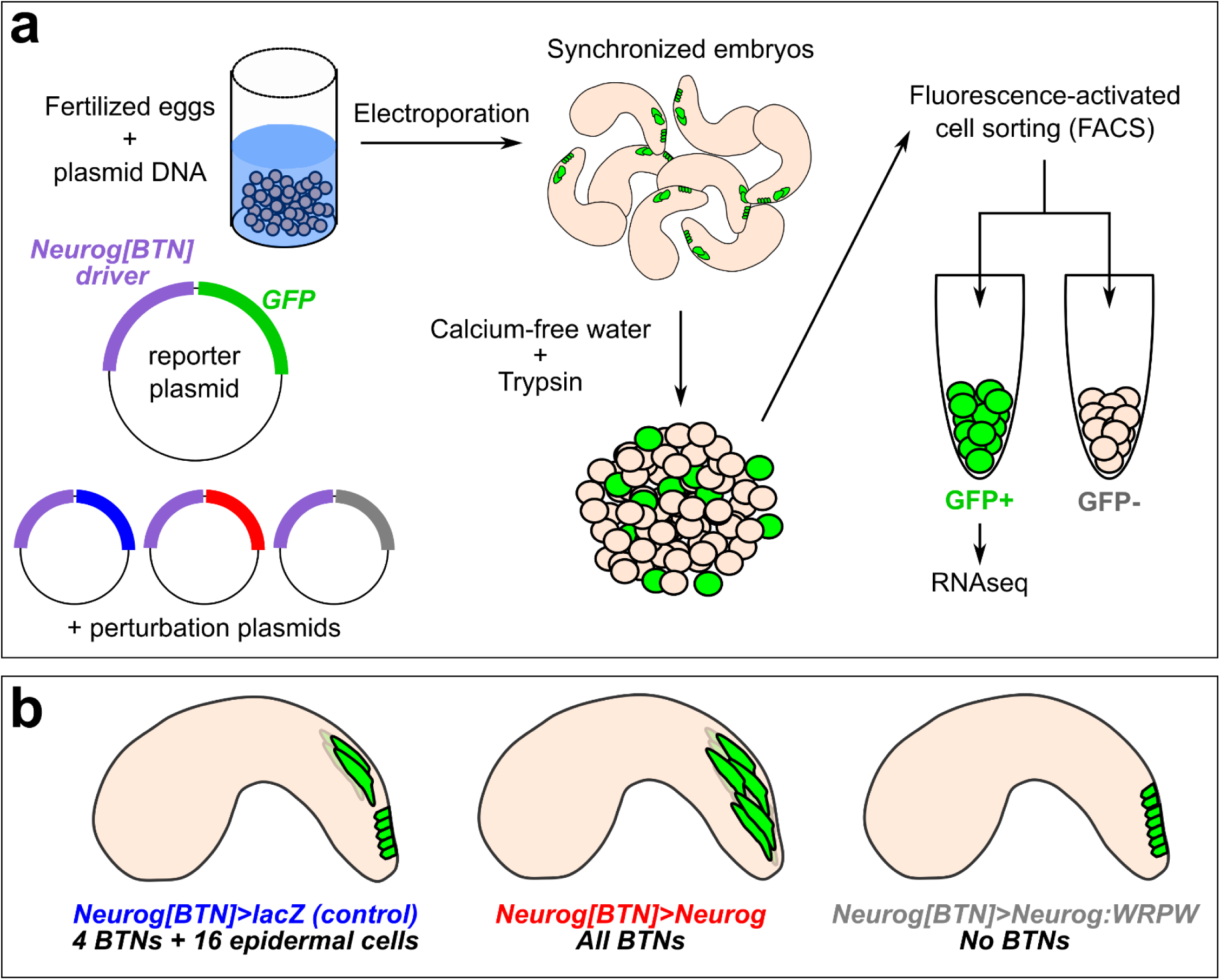
FACS and RNAseq methodology a) Schematic of FACS-RNAseq used in this study. b) Schematic of different conditions used to sort “control” BTN lineages and Neurog gain- or loss-of-function.

Under these conditions, 522 target genes were upregulated by Neurog (LogFC>0.6, p<0.05) and 176 downregulated by Neurog::WRPW (LogFC<-0.6, p<0.05), with 76 genes in both categories (**Fig 3a**). The larger number of genes upregulated by Neurog was expected, given that many more ectopic BTNs are specified in *Neurog>Neurog* than the number of BTNs lost in *Neurog>Neurog::WRPW (Stolfi et al., 2015)*. However, this could also be an artefact due to lower statistical support as a result of vastly different numbers of cells sorted between *Neurog>Neurog::WRPW* replicates (2418 cells and 114 cells). Although there were reported whole-mount *in situ* hybridization (ISH) images for 33 of these 76 genes on the ANISEED tunicate expression database (Brozovic et al., 2018), we were able to infer clear BTN expression from such database images for only 10 genes. These included the marker gene *Asic* previously used to assay BTN specification (Coric et al., 2008), and additional genes such as *alpha-Tubulin* (KH.C8.892), *Rgs19/20 (KH.C1.314), Slc35g2 (KH.L141.43), Bassoon-like (KH.C5.481), Onecut*, and others with no substantial homology to known proteins. Because several other known BTN markers were not represented, we relaxed our criteria. More specifically, we looked at genes that were upregulated by Neurog (1444) and downregulated by Neurog::WRPW (1303) with no p-value cutoff. This increased the overlapping set, and thus our candidate target gene list, to 372 genes (**Fig 3b**).

**Figure 3.**
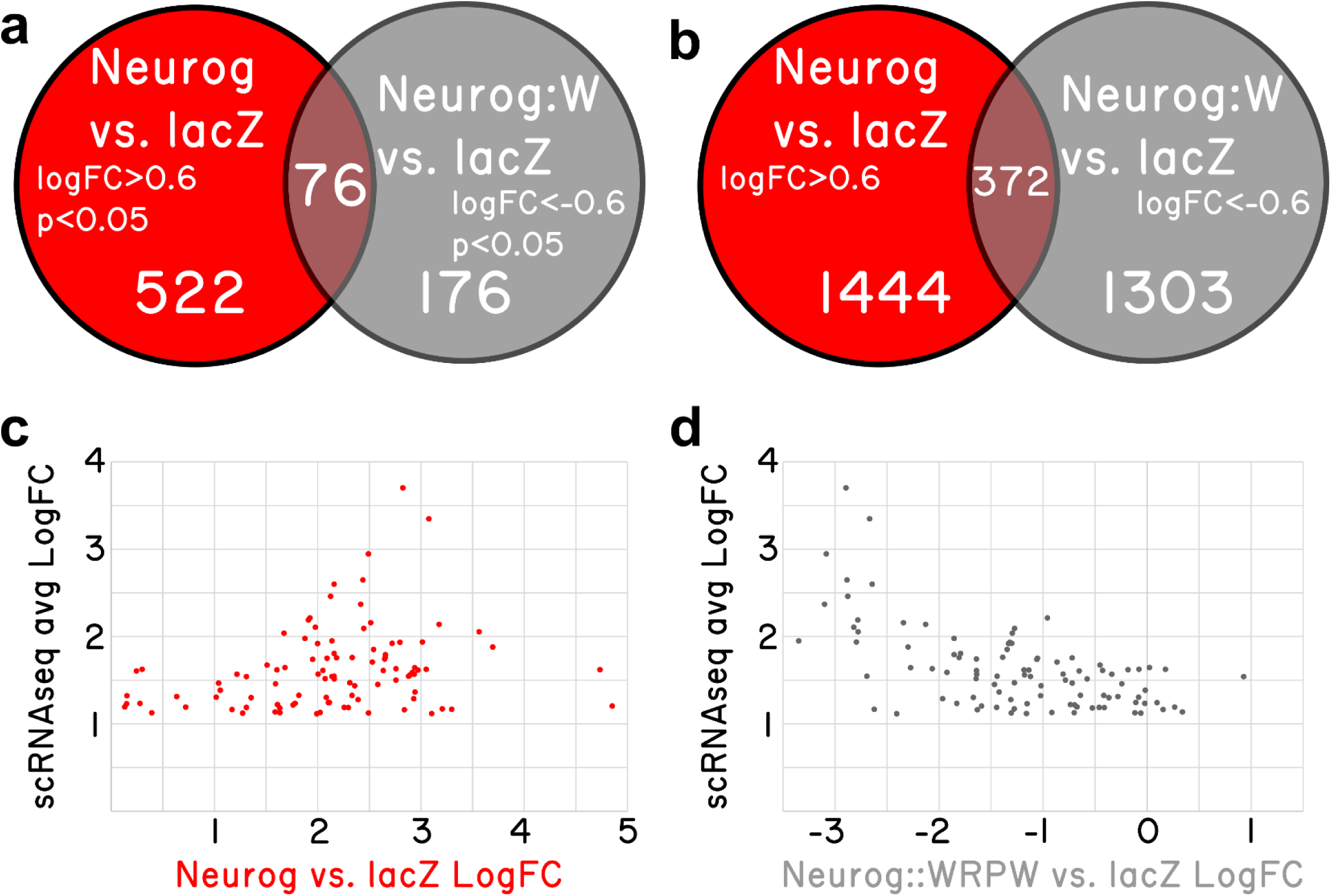
RNAseq-based analyses of Neurog function in BTNs. a) Non-proportional Venn diagram indicating number of genes in each condition showing statistically-significant (p<0.05) differential expression (LogFC>0.6 in Neurog vs. lacZ or <-0.6 in Neurog::WRPW vs. lacZ). b) Same analysis as in (a) but with p-value cutoff removed. c) Comparison of “avg LogFC” of top 98 BTN genes identified by single-cell RNAseq (Horie et al., 2018) to LogFC in Neurog vs lacZ, showing that all 98 are positively upregulated by Neurog. d) Similar comparison as in (c) but to LogFC in Neurog::WRPW vs. lacZ, revealing that all but 7 of the top 98 BTN genes are downregulated by Neurog::WRPW.

To see if we were measuring meaningful expression of Neurog targets in the BTNs. We crossreferenced this differential gene expression data to a previously published single-cell RNAseq data set comprising the top 100 genes enriched in the BTNs relative to other cell types at 12 hpf at 18°C (Horie et al., 2018)(**Supplemental Table 2**), with the exception of two genes: *KH.S1555.2* (which was not present in our dataset) and *Neurog* (due to confounding reads from the electroporated plasmids). We found that all 98 top BTN genes in the scRNAseq dataset were positively regulated by Neurog overexpression (LogFC>0, **Fig 3c**). Similarly, 91 of 98 top genes were negatively regulated by Neurog::WRPW overexpression (LogFC<0, **Fig 3d**). This confirmed that Neurog positively regulates BTN fate, and that our strategy was able to detect differential Neurog target gene expression in the BTNs, though statistical support might be lacking for many BTN markers at the embryonic stages that were sequenced.

### Validation of BTN genes by *in situ* hybridization

Because the above results suggested our differential expression analysis criteria might 1) be too stringent to detect all real Neurog targets in the BTNs and 2) might contain false-positives associated with leaky expression of the Neurog driver in other tissues, we decided to validate a large set of potential BTN markers by fluorescent ISH (**Supplemental Table 3**). We successfully prepared probes for 137 genes, from a mixture of cDNA clones, RT-PCR, and synthetic DNA templates (see **Methods** for details, and **Supplemental Table 3** for all probe template sequences). Of these, 49 were confirmed to be upregulated in the migrating BTNs (**Fig 4**). For another 29, it was not clear if they were expressed in BTNs or not, due to low signal or obscuring signal from neighboring tissues. Most are likely true positives, but confirming them will require better probes or higher resolution imaging. 15 genes showed CNS-specific expression, but in other neurons, 15 showed expression mainly in non-neural tissues, and 29 were true “negatives” with no or little signal throughout the whole embryo (data available upon request).

**Figure 4.**
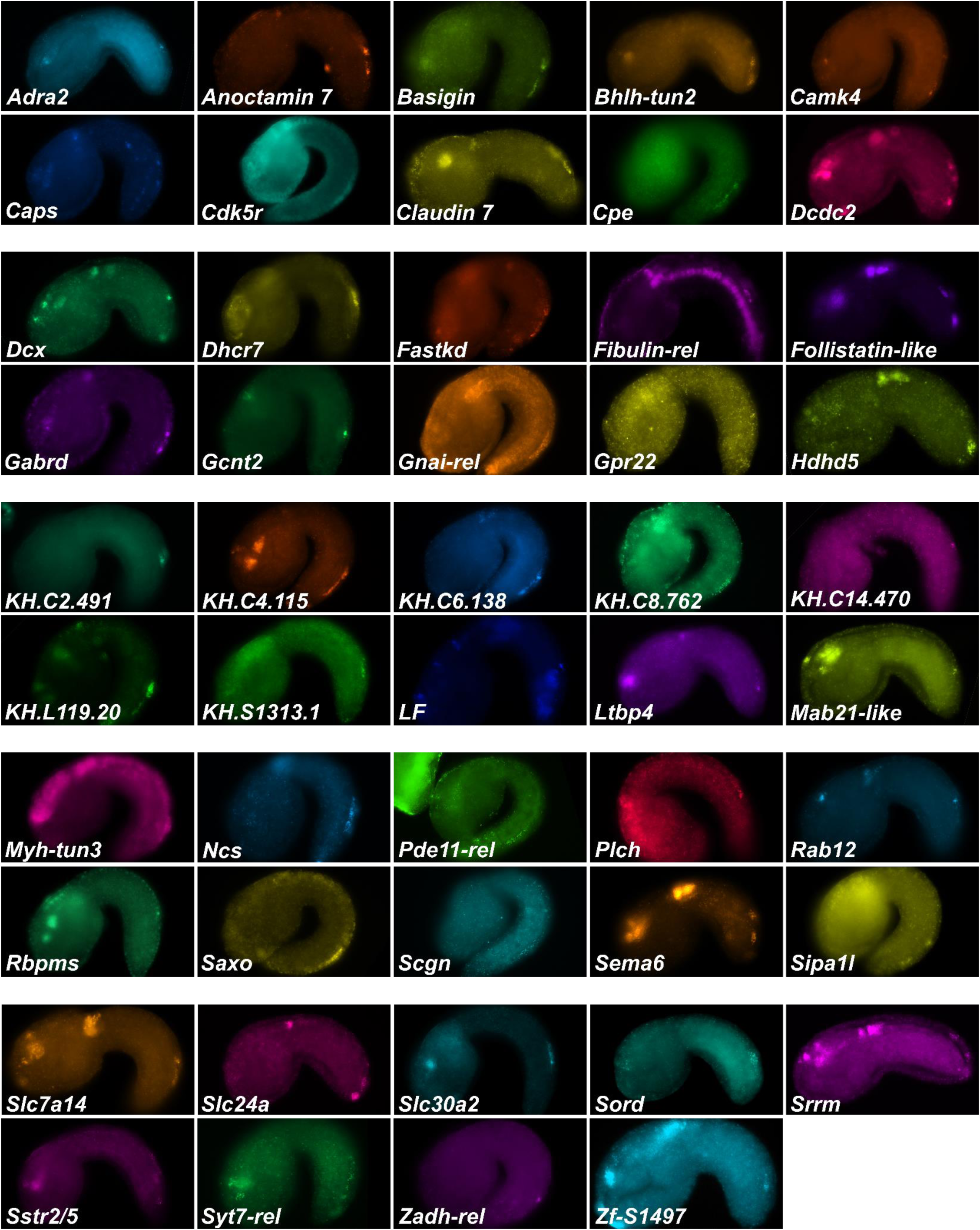
Fluorescent whole-mount *in situ* hybridization of Neurog targets (all 49 validated candidates).

From our results it became obvious that validation of BTN expression by ISH in this subset correlated most closely with overall transcript abundance in the samples. 22 of the top 50 genes with highest LogCPM were BTN+, with another 10 showing “unclear” signal. In contrast, only 3 of the 50 genes with <u>lowest</u> LogCPM were BTN+, though 11 were “unclear”. 23 of the bottom 50 genes were “negative”, suggesting that many of these might in fact be expressed in the BTNs, but at levels that are below the threshold of detection by ISH. Among those genes that were validated by ISH as specifically upregulated in BTNs during delamination and axon extension, some are expressed in either the aBTN or pBTN alone, though it is unclear if this indicates merely a difference in timing of gene expression between the two. However, there is reason to believe that there are functional differences between the aBTN and pBTN. For instance, the gabaergic marker *Glutamate decarboxylase (GAD)(Zega et al., 2008)* is only ever seen to be expressed in the aBTN (**Fig 1c**), while the cholinergic marker *Slc18a3/Vesicular acetylcholine transporter (Vacht)(Yoshida et al, 2004)* can be seen in the pBTN (**Fig 1b**). Both are upregulated by Neurog (*GAD* LogFC = 2.7, *Slc18a3* LogFC = 3) and downregulated by Neurog::WRPW (*GAD* LogFC = −1.4, *Slc18a3* LogFC = −1.1), suggesting that *Neurog* might regulate both targets but in separate aBTN/pBTN contexts.

We also found that many genes were expressed in other CNS neurons in addition to BTNs. Such genes are likely to be targets of Neurog in these other CNS neurons, especially in the Motor Ganglion (MG) and brain. Thus, Neurog is likely to regulate overlapping sets of genes that can be broadly neuronal, BTN-specific, or aBTN/pBTN-specific, highlighting the importance of combinatorial regulation with other lineage-specific transcription factors is regulating neuronal subtype-specific fates and gene expression.

### Predicting BTN effector gene functions

Several candidate Neurog-targets in the BTNs appear to be involved in neuronal function, especially neurotransmission, suggesting relatively early transcription of such genes relative to larval hatching. These include *GABA receptor (Gabrd), Anoctamin 7 (Ano7), Nenronal calcium, sensor (Ncs), Adrenergic receptor alpha 2 (Adra2), Synaptotagmin 7-related (Syt7-rel)*, the neuropeptide-encoding *Ci-LF precursor (LF) (Kawada et al., 2011)*, and others (**Fig 4**). Due to our interest in understanding the delamination, migration, and dynamically polarized axon outgrowth of the BTNs, we focused our analysis on those genes hypothesized to be more directly involved in such cell behaviors. These encompassed both intracellular and extracellular proteins, and their proposed functions have begun to paint a picture of the mechanisms underlying BTN morphogenesis.

#### Cdk5 regulatory subunit (Cdk5r) and Doublecortin (Dcx)

Microtubule stabilization has been shown to be essential for neuronal migration and axon specification (Witte et al., 2008), though the mechanisms underlying its local control remain largely unknown (Kapitein and Hoogenraad, 2015). In vertebrates, Neurog1 and Neurog2 control neuronal migration in part through upregulation of *Cdk5r1* and *Doublecortin (Dcx)* effectors (Ge et al., 2006). Both *Ciona* orthologs of *Cdk5r1* and *Dcx* are upregulated in BTNs by Neurog (**Fig 5a**)(Horie et al., 2018), suggesting a conserved regulatory network for neuronal migration that is shared between *Ciona* and vertebrates. Cdk5r1 (also known as p35) is an activator of Cdk5, and the Cdk5/Cdk5r1 is required for microtubule stability in neuronal migration and axon outgrowth in several examples (Chae et al., 1997; Lambert de Rouvroit and Goffinet, 2001; Nikolic et al., 1996; Smith et al., 2001). Human DCX and the closely related Doublecortin-like kinases (DCLK1/2) are represented by a single ortholog in *Ciona, Dcx/Dclk* (referred from here on as simply *Dcx*). In mammals, Dcx has been proposed to be essential for neuronal migration and differentiation by nucleating, binding, and/or stabilizing microtubules (Corbo et al., 2002; Ettinger et al., 2016; Moores et al., 2004). The closely related vertebrate Doublecortin-like kinases are also associated with microtubules (Lin et al., 2000). While Dclk1 mutant mice show few neuronal migration defects, Dclk1/Dcx double mutants show extensive cortical layering and axonal defects, suggesting some overlapping roles for these paralogs (Deuel et al., 2006). Dcx/Dclk proteins contain two DCX protein domains, as does *Ciona* Dcx. As a proxy for the subcellular localization of this protein, we constructed a DcxΔC::GFP fusion comprised of the two DCX domains fused to GFP. When driven by the *Ebf* neuronal promoter (−2.6kb upstream)(Stolfi and Levine, 2011) in differentiating neurons, we observed DcxΔC::GFP enrichment in microtubule bundles extending into the axons of BTNs (**Fig 5b**). This microtubule bundle localization suggests a conserved role for Dcx in *Ciona*.

**Figure 5.**
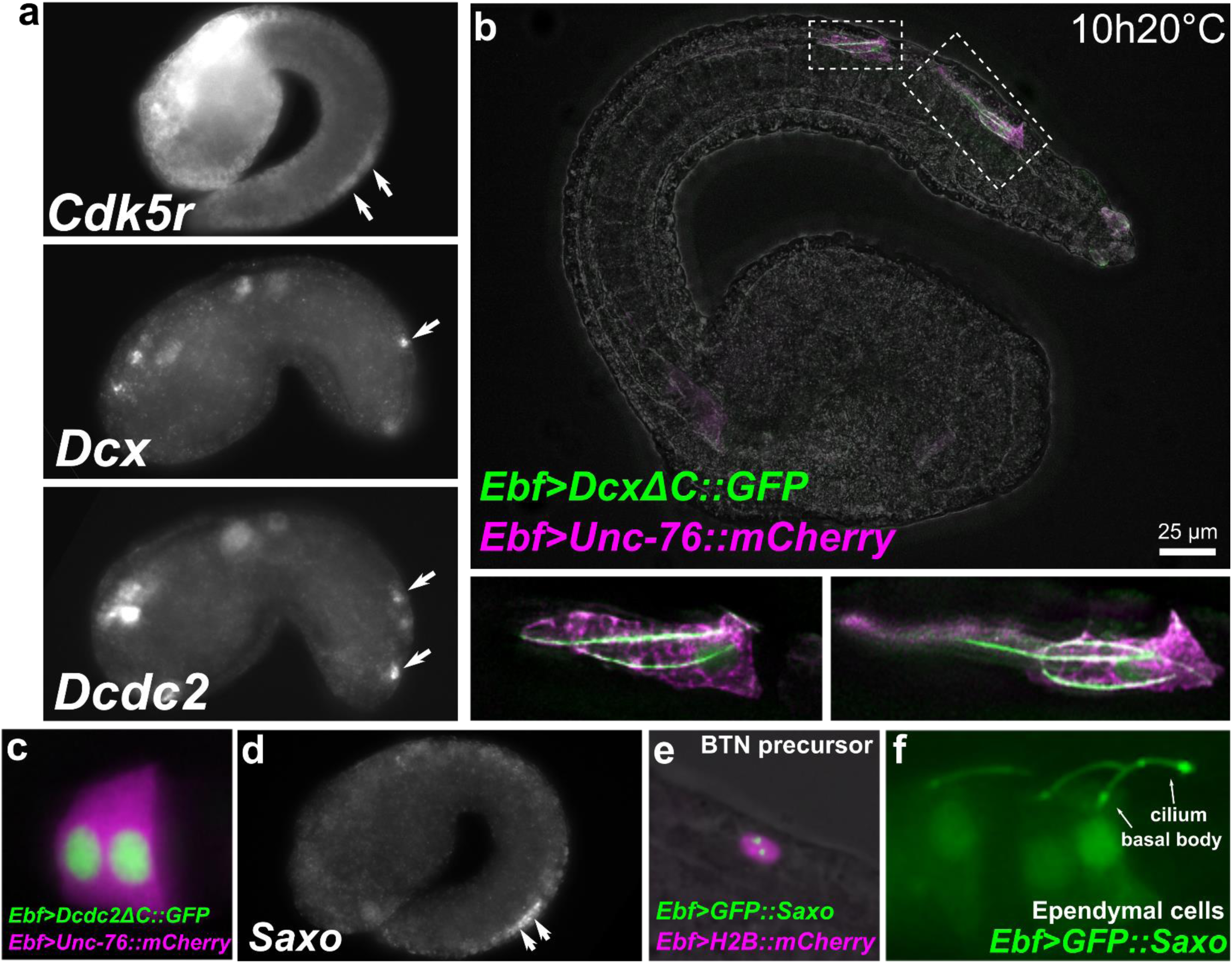
Candidate effectors of microtubule dynamics expressed in BTNs. a) *In situ* hybridization of *Cdk5r, Dcx*, and *Dcdc2*. Arrows indicate BTNs. b) Embryo electroporated with *Ebf>DcxΔC::GFP* and *Ebf>Unc-76::mCherry* plasmids, showing GFP labeling of microtubule bundles in migrating BTNs (insets of fluorescence channels magnified at bottom). c) Embryo electroporated with *Ebf>Dcdc2ΔC::GFP* and *Ebf>Unc-76::mCherry* plasmids, showing that despite having 2 DCX domains like Dcx, Dcdc2 is localized mostly to the nucleus, like its mammalian ortholog (Coquelle et al., 2006). d) *In situ* hybridization of *Saxo*, showing BTN-specific expression. Expression was also seen in ddNs as reported in (Gibboney et al., 2019), which are out of focus. e) GFP::Saxo labeling putative centrioles/centrosome in a BTN precursor at mid-tailbud stage. f) GFP::Saxo was also seen in the basal bodies and associated cilia in ciliated ependymal cells of the neural tube/nerve cord.

Another DCX-family member was found to be upregulated in BTNs, namely *Doublecortin domain-containing 2 (Dcdc2*, **Fig 5a**). This gene also encodes for a protein that contains two DCX domain. However, when we fused the DCX domains from Dcdc2 to GFP (Dcdc2ΔC::GFP), we observed highly specific nuclear localization (**Fig 5c**), in stark contrast to the microtubule-associated localization of DcxΔC::GFP. Nuclear localization of human DCDC2 has been reported previously (Coquelle et al., 2006), further confirming the likely diverse functions of DCX domains in localizing DCX-family members to very distinct subcellular compartments.

#### Saxo: Stabilizer of axonemal microtubules

Positioning of the centrosome and associated Golgi apparatus has been shown to be an essential intrinsic cue for neuronal polarization (Andersen and Halloran, 2012; de Anda et al., 2010). However, this appears to be highly context-dependent and difficult to study *in vivo* due to the transient nature of centrosome position, tissue complexity in the developing CNS, and species- and cell-type-specific differences (Basto et al., 2006). Microtubule stabilization has been shown to be essential for axon specification (Witte et al., 2008), though the mechanisms underlying its local control remain largely unknown (Kapitein and Hoogenraad, 2015). Because centrosome repositioning is also driven by microtubule stabilization (Burute et al., 2017; Pitaval et al., 2017), this suggests that such centrosome-associated microtubule stabilizers might function as key effectors linking centrosome position and axon outgrowth. In the BTNs, initial axon outgrowth is concurrent with migration: the leading edge of the BTNs extends and becomes the proximal (anterior) process of the axon. Thus, polarization, migration, and axon outgrowth might be tightly coupled in the BTNs

Previous MG neuron transcriptome profiling and a follow up ISH revealed that *Saxo (Stabilizer of axonemal microtubules)* was expressed in the BTNs (**Fig 5d**), in addition to the ddNs (Gibboney et al., 2019). *Saxo* is the *Ciona* ortholog of human *SAXO1/SAXO2*, formerly *FAM154A/FAM154B*. These genes encode a highly conserved subfamily of STOP/MAP6-related proteins that stabilize microtubules (Dacheux et al., 2015). In human cell culture, SAXO1 localizes to centrioles and cilia and mediates stabilization of cold-resistant microtubules. They do so through 7 microtubule-binding/stabilizing “Mn” domains (Dacheux et al., 2015), which are conserved in *Ciona Saxo*. SAXO1/2 have not been implicated in neurodevelopment or cell polarity *in vivo*, but depletion of related MAP6 proteins in mice results in synaptic defects and schizophrenia-like symptoms (Volle et al., 2012).

A GFP::Saxo fusion when expressed in *Ciona* was found to localize to centrosomes in BTN precursors (**Fig 5e**), and to cilia of ependymal cells (**Fig 5f**), also consistent with a potentially conserved role in microtubule stabilization. Given its expression in both BTNs and ddNs, and given the dynamic repositioning of the Golgi apparatus observed in both these neurons types immediately predicting direction of axon outgrowth (Gibboney et al., 2019; Stolfi et al., 2015), Saxo is one of the more intriguing candidate effectors of neuronal polarization that remain to be functionally characterized.

How might extracellular cues impinge on centrosome position *in vivo?* One pathway that has been implicated in this process during neuronal migration is the Semaphorin/Plexin pathway (Renaud et al., 2008). We found that *Semaphorin 6 (Sema6)*, a class 6 Semaphorin orthologous to human SEMA6A/SEMA6B/SEMA6C (Yazdani and Terman, 2006) is expressed in migrating BTNs and broadly in other CNS neurons including those in the brain and MG (**Fig 4**). In mice, *Sema6a* and its receptor *Plexin A2* control migration in granule cells of the cerebellum, through regulating centrosome position and nucleokinesis (Renaud et al., 2008). In mammals, Sema6a can inhibit Plexin in *cis* as a mechanism to reduce sensitivity to Sema6a in *trans (Haklai-Topper et al., 2010)*. Perhaps its expression in developing *Ciona* larval neurons reflects such a mechanism.

### TGFBeta inhibitors expressed by the BTNs

Among the predicted *extracellular* proteins that identified as Neurog targets in the BTNs/DDNs, genes encoding extracellular regulators of TGFβ signaling stood out, namely *Latent TGFβ-binding protein 4 (Ltbp4), Follistatin-like (Fstl)*, and *Fibulin-related* (**Fig 6a**). All encode ECM components that bind TGFβ to modulate its activity (Chang, 2016; Robertson and Rifkin, 2016). TGFβ is the main cue that induces axon formation in the mammalian brain (Jason et al., 2010). The *Ciona* ortholog of *TGFβ1/2/3 (TGFβ, KH.C3.724)* is expressed in the notochord, and *Type I TGFβ Receptor (KH.L22.40)* is expressed in the CNS including BTNs (Imai et al., 2004). *TGFβ* expression in the notochord itself is polarized, becoming more and more restricted to the posterior tip of the notochord later in development (Reeves et al., 2017). Thus, *Ciona* notochord-derived TGFβ may function as a guidance cue for the dynamic polarization of migrating BTNs, which invert their polarity to extend the distal axon branch towards the tail tip precisely when *TGFβ* expression shifts posteriorly. However, the expression of TGFβ-interacting proteins by the BTNs themselves might also be crucial for TGFβ-dependent polarization. More specifically, we hypothesize that deposition of Ltbp4, Fstl, and/or Fibulin by migrating BTNs might sequester TGFβ ligand along its migration route, promoting continued forward migration/axon outgrowth at first. Later, when *de novo TGFβ* transcription is maintained in the posterior tip of the notochord, newly synthesized TGFβ might overcome this temporary inhibition, and shift the balance of TGFβ signaling posteriorly, thus inverting the polarity of axon outgrowth by the BTNs (model summarized in **Fig 6b**). In this model, both intrinsic and extrinsic sources of extracellular effectors are required to direct BTN polarization.

**Figure 6.**
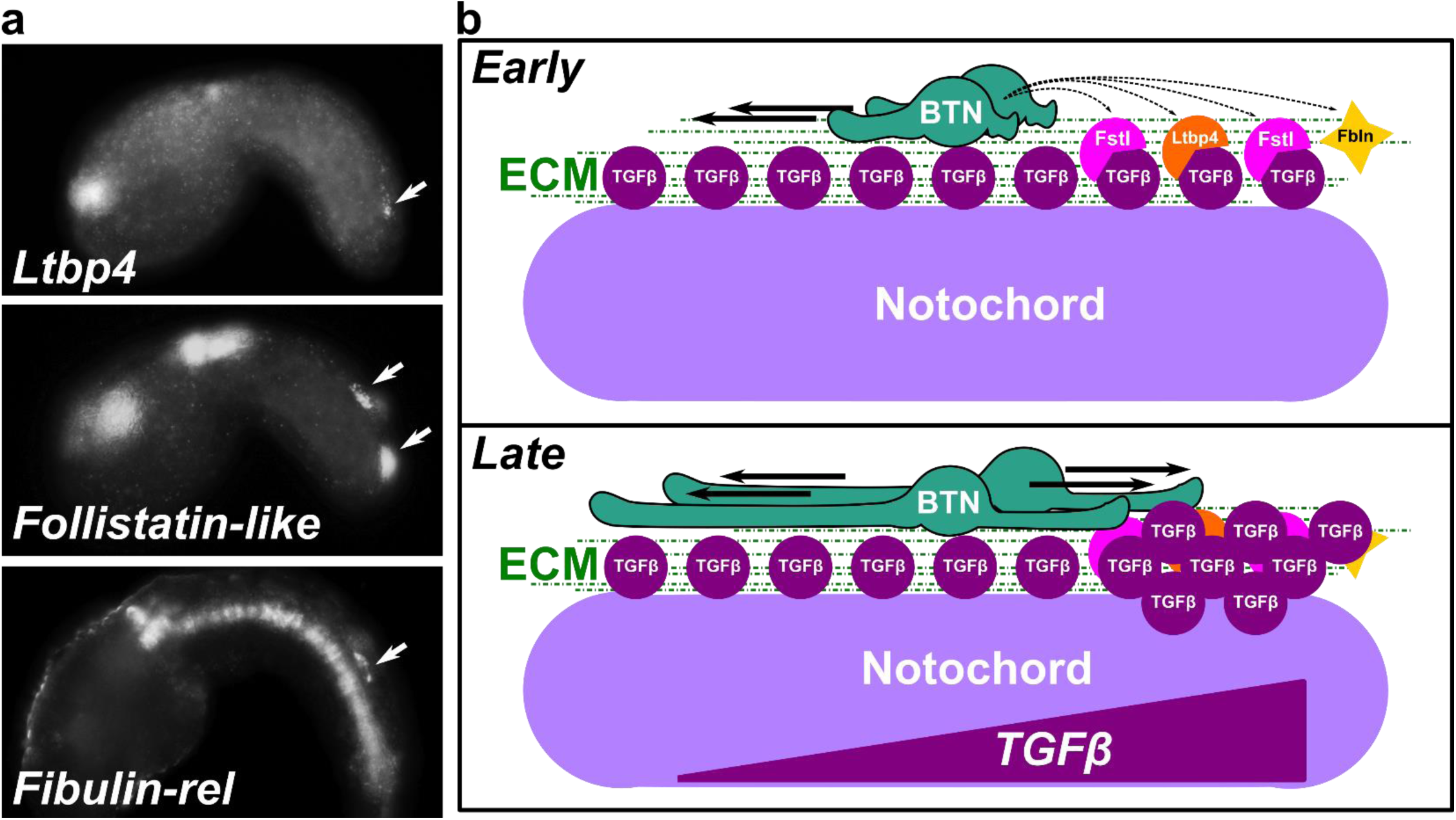
Antagonists of TGFβ signaling expressed in migration BTNs and model for TGFβ-based BTN polarization. a) *In situ* hybridization for genes encoding secreted antagonists of TGFβ ligands (*Ltbp4* and *Follistatin-like*) and of TGFβ receptors (*Fibulin-related*). Arrows indicate BTNs. b) In our proposed (but untested) model for BTN polarization, the notochord is a source of TGFβ ligand, which is essential for BTN axon outgrowth. At earlier tailbud stages, migrating BTNs secrete TGFβ signaling inhibitors (Fstl, Ltbp4, Fbln) as they move forward. This polarized inhibition ensures forward axon extension. Later in development, when restricted TGFβ expression in posterior notochord reaches its peak (Reeves et al., 2017), new TGFβ molecules overcome local inhibition imposed by BTN migration path. BTN axons are now equally attracted anteriorly and posteriorly, resulting in two axon branches and characteristic bipolar morphology.

#### Gai-related

We identified a gene encoding a homolog of the small Gαi/o protein subunit family that by *in situ* hybridization was observed to be upregulated in migrating BTNs and notochord cells (**Fig 7a**)(Reeves et al., 2017). This rather divergent *Gαi* gene (*KH.C2.872*, referred to simply as *Gnai-related*, or *Gnai-rel*), is one of three *Gαi/o* paralogs that seem to be *Ciona*- (or tunicate-) specific duplications: *KH.C1.612, KH.C2.872*, and *KH.L96.27*. Of these, *KH.C1.612* seems to be the original “founding” paralog, as it still retains exons/introns, while *KH.C2.872*, and *KH.L96.27* are both encoded by a single exon, suggesting possible duplication by retrotransposition, followed by subfunctionalization (Ohno, 2013).

**Figure 7.**
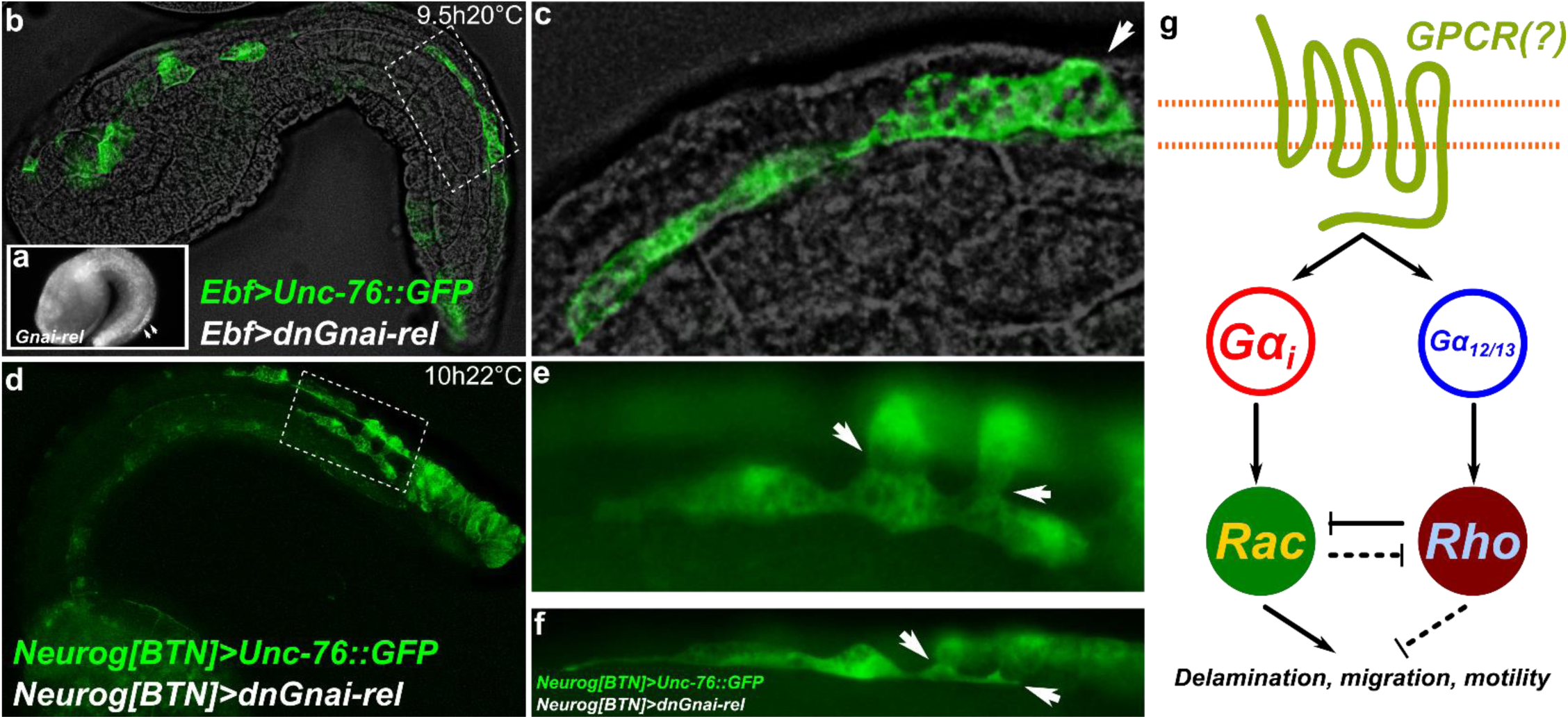
Gαi function is required for proper delamination of BTNs. a) Inset showing BTN- and notochord-specific expression of *Gnai-related*. Arrowheads indicate BTNs. b) Embryo electroporated with *Ebf>Unc-76::GFP* (green) and *Ebf>dnGnai-rel S48N*. Boxed area magnified in c) showing failure of BTN to completely separate from epidermal surface (arrowhead), d) Embryo electroporated with *Neurog[BTN]>Unc-76::GFP* (green) and *Neurog[BTN]>dnGnai-rel S48N*. Single focal plane corresponding to boxed area magnified in e) showing prolonged adhesion between BTN and epidermal cells (arrowheads), f) Another embryos showing similar cell-cell adhesions (arrowheads), which should be completely shed by this stage. g) Proposed model (based on mechanisms found in other cell types and organisms) for developmental “switch” based on opposing functions of Gαi and Gα12/13. Dashed lines indicated possible reciprocal and/or redundant mechanisms.

The developmentally regulated *Gnai-rel* expression pattern was intriguing, and suggestive of a rate-limiting role for this gene in cellular movements, since it is specifically upregulated in migrating BTNs and intercalating notochord cells. To test its possible function in BTN cell behaviors, we overexpressed a dominant-negative form of Gnai-rel (S48N)(Barren and Artemyev, 2007), using either the *Ebf or Neurog[BTN]* drivers. In both cases, BTNs failed to delaminate from the dorsal epidermis midline, although the cells still appeared to differentiate and attempt to migrate and extend their axons forward (**Fig 7b-f**). These results suggest that upregulation of *Gnai-rel* by Neurog is necessary for proper BTN delamination.

In mammalian cells, upregulation of Gαi can act as a molecular “switch” to inhibit RhoA by competing with Gα12/13 proteins for interactions with the same G-protein coupled receptor (GPCR), resulting in the activation of Rac1 activation and increased cell motility(Sugimoto et al., 2003). This antagonism between Rho/Rac is also seen in delaminating neural crest cells, in which Rho inhibits Rac activity to keep cells in an epithelial state (Shoval and Kalcheim, 2012). In radial neuron migration, Gα12/13 proteins *terminate* migration(Moers et al., 2008), and have been shown to do so through RhoA in cultured neurospheres(Iguchi et al., 2008). Thus, transcriptional control over the relative expression levels of Gαi and Gα12/13 might be a common mechanism for regulation of neuronal migration, shifting between activation of Rac1 (promoting migration) or of RhoA (inhibiting migration)(**Fig 7g**). Whether *Gnai-rel* has been further subfunctionalized to interact specifically with a given GPCR, or if its function is simply to provide a tissue-specific “boost” of Gαi at different points in development, is an evolutionarily interesting question that remains to be answered.

### Conclusions

Here we report the transcriptional dynamics of effector genes downstream of Neurog in the BTNs. We have identified and validated several predicted BTN-specific genes that might be key for BTN cell behaviors, especially their dynamic but stereotyped delamination, migration, and polarization processes. We have identified one gene, *Gnai-rel*, as an effector gene whose activity is necessary for BTN delamination, potentially through a conserved, chordate-specific “switch” for Rho/Rac activity.

## Methods

### FACS and RNAseq

Embryos were electroporated with the following combinations of plasmids: *70 μg Neurog −3010/−773+-600>tagRFP/tagBFP* + *50 ug Neurog −3010/−773+-600stop>Neurog (Neurog>Neurog* condition). *70 μg Neurog −3010/−773+-600>tagRFP/tagBFP* + *50 ug Neurog −3010/−773+-600stop>Neurog::WRPW (Neurog>Neurog::WRPW* condition), *70 μg Neurog −3010/−773+-600>tagRFP/tagBFP* + *50 ug Neurog −3010/−773+-600>lacZ (Neurog>lacZ* “control” condition). Embryos were dissociated and FACS-isolated using a BD FACS Aria cell sorter into lysis buffer from the RNAqueous-Micro RNA extraction kit (ThermoFisher) as previously established (Wang et al., 2018a, b). BFP+ or RFP+ cells were isolated with no counterselection. Cell numbers obtained were: Neurog>lacZ(control) replicate 1: 975 cells; Neurog>lacZ(control) replicate 2: 200 cells; Neurog>Neurog replicate 1: 284 cells; Neurog>Neurog replicate 2: 800 cells; Neurog>Neurog::WRPW replicate 1: 2418 cells; Neurog>Neurog::WRPW replicate 2: 114 cells. RNA was extracted from each sample according to the RNAqueous-Micro kit instructions. cDNA synthesis was performed as described (Wang et al., 2017), with SMART-Seq v4 Ultra Low Input RNA kit (Takara). Sequencing libraries were prepared as described (Wang et al., 2017), with Ovation Ultralow System V2 (NuGen). Libraries were pooled and sequenced by Illumina NextSeq 500 Mid output 150 Cycle v2, to generate 75 bp paired-end reads, resulting in 192,396,840 single-end reads for the 6 samples. Resulting FASTQ files were processed by STAR 2.5.2b and mapped to the C. *robusta* genome (Dehal et al., 2002; Satou et al., 2008). Output bam files were processed using Rsubread/featureCounts (Liao et al., 2013), with the parameter “ignoreDup=TRUE” to remove the read duplications resulting from library amplification. All reads after duplication removal that mapped to the exons of KyotoHoya (KH) gene models (Satou et al., 2008) were counted for differential expression analysis. Differential expression beween Neurog>Neurog and Neurog>lacZ, and between Neurog>Neurog::WRPW and Neurog>lacZ was measured by EdgeR (Robinson et al., 2010)(**Supplemental Table 1**). Raw data, counts, and scripts can be found at https://osf.io/uqfn2/

### Embryo *in situ* hybridizations

Adult *Ciona robusta* (*intestinalis* Type A) were collected from San Diego, CA (M-REP). Dechorionated embryos were obtained and electroporated as previously established (Christiaen et al., 2009a, b). Sequences of *in situ* hybridization probe templates can be found in **Supplemental Table 3**. *Neurog* perturbation and control plasmids were previously published (Stolfi et al., 2015). Probes were prepared either from published clones, synthetic DNA fragments (Twist Bioscience), or directly from RT-PCR amplicons (see **Supplemental Table 3** for details). Probe synthesis and fluorescent, whole-mount *in situ* hybridization were carried out as previously described (Beh et al., 2007; Ikuta and Saiga, 2007). Images were captured using Leica DMI8 or DMIL LED inverted epifluorescence compound microscopes. Plasmid sequences not previously published can be found in the Supplemental Sequences file hosted at https://osf.io/uqfn2/

## Acknowledgments

We thank members of our lab at Georgia Tech for their thoughtful feedback and technical assistance throughout this project. We thank Wendy Reeves and Michael Veeman for the *Gnai-related* probe template. This work is funded by NIH award R00 HD084814 to A.S.

